# Pushing the limits of SCP: bacSCP, a proof-of-concept study to investigate heterogeneity of bacteria by single cell proteomics

**DOI:** 10.64898/2026.04.09.717485

**Authors:** Julia Leodolter, Tim Thierer, Karl Mechtler, Manuel Matzinger

## Abstract

Single-cell proteomics (SCP) has emerged as a powerful approach to quantify protein expression variability at cellular resolution, yet most state-of-the-art workflows are tailored to eukaryotic cells with only one study exploring how single bacteria can be analyzed by mass spectrometry. Here, we established bacSCP, a protocol extending SCP to bacterial cells, facing analytical challenges such as the thick bacterial cell wall hampering lysis, the extremely small cell size and resultant low protein content, and the consequently relatively high level of contaminating proteins from external sources. Using this bacSCP pipeline, we quantified more than 50 bacterial proteins from single *Bacillus subtilis* and *Escherichia coli* cells. Upon heat stress, we reproducibly observed up to 8-fold upregulation of chaperones including GroEL, GroES, and ClpC for a *B. subtilis ΔmcsB* strain. Importantly, single-cell measurements revealed potential heterogeneity within the heat-stressed subpopulation, enabling interrogation of stress-response variability at the proteome level. These results demonstrate the feasibility of bacSCP and provide a foundation for studying bacterial stress adaptation and phenotypic diversity with single-cell proteomic resolution.

## INTRODUCTION

Single Cell Proteomics (SCP) by mass spectrometry (MS) is an emerging technology that continuously pushes its limits towards improved reproducibility, throughput and sensitivity. As the protein content of a single eukaryotic cell is very low in the 200 pg range, SCP requires state-of-the-art equipment and workflows to reduce sample loss in an extremely low input setting. Successful approaches apply semi-automated sample preparation within minimal volumes using the cellenONE, which performs ultra-precise image-based cell sorting and droplet dispension. Downstream processing of cells in a One-Pot reaction setup was shown to minimize losses^1–3^. Using such lossless workflows and the Astral mass analyzer achieved remarkable coverage of more than 5000 proteins quantified on average per single cell. Such workflows were applied to investigate heterogeneity based on different cell-cycle phases even from cultured untreated cells and to precisely follow cell differentiation and map it to RNAseq data from large cohorts of thousands of cells^4–6^.

While eukaryotic SCP is already a challenging task necessitating the most advanced hardware, downscaling this technique to bacteria pushes the boundaries of current technology. With an expected volume of ∼2 fL^7^, bacteria are approximately 1000 times smaller than eukaryotic cells (∼2 pL)^8^. and thus contain much less protein, with bacterial cell content estimated to be below 1 pg protein even for actively growing cells^9^.

Major challenges with bacteria include efficient cell sorting due to small cell size and sample loss during handling due to low input material. Additionally, cell lysis needs to be compatible with downstream MS applications, while effectively breaking the bacterial cell wall. Gram-positive bacteria (e.g. *Bacillus subtilis*) are surrounded by a thick peptidoglycan cell wall, while Gram-negative bacteria (e.g. *Escherichia coli*) have a peptidoglycan wall, a periplasmic compartment and an outer membrane. To the best of the authors’ knowledge there is currently only a single publication that successfully attempted first steps towards single bacteria proteomics, reporting 12 confident proteins from single bacteria using a modified SCoPE MS approach, where they multiplexed 9 cells plus an additional 250 cell carrier to push sensitivity^10^. Although this sensitivity boost was necessary with the hardware available at the time of publication, such DDA-TMT based approaches suffer from a bias towards quantifying peptides present in the carrier proteome, which hampers potential detection of new or rare cellular subpopulations, as well as from limited dynamic range and ratio compression^3^.

Expanding on these results and establishing robust bacterial SCP will be critical to enable direct proteomic studies of bacterial heterogeneity at the single-cell level. Phenotypic heterogeneity in isoclonal bacterial populations exists as a bet-hedging strategy, where individual bacteria stochastically shift to different metabolic states, which allows rapid adaptation to environmental changes, such as nutrient and temperature stresses^11,12^. Importantly, bacterial population heterogeneity plays a crucial role in persister cell formation during antibiotic stress and during infection^13^.

Analysis of bacterial heterogeneity at the single cell level is already well established using fluorescence-based phenotypic observations of cellular processes, as well as through transcriptomics and metabolomics ^12,14^. Integrating SCP with these existing approaches would provide a critical layer of information for understanding bacterial heterogeneity, as cellular regulation occurs not only on DNA and RNA, but also on protein level, through post-translational modifications, targeted degradation and sequestration of key factors, especially during cellular stress^15,16^.

In bacteria, heat stress is a well-studied environmental challenge. During heat stress the protein quality control machinery is upregulated and critical to maintain protein homeostasis and to protect cells from proteotoxic stress. The universal essential folding chaperonin GroEL-GroES refolds damaged proteins, while proteases such as the ClpCP system degrade damaged proteins^17^. In this study, we establish a label-free proof-of-concept approach for bacterial single cell proteomics (bacSCP) for the first time. Thereby we adapt and expand current methods available for eukaryotic and bacterial SCP. We benchmark methods for bacterial cell isolation and cell opening to allow for protein extraction and digestion. When we apply our method to heat stressed *B. subtilis ΔmcsB* strain, we detect upregulation of the critical heat shock response proteins ClpC, GroES and GroEL and investigate the possibility to distinguish heat-stressed from non-stressed cells at single cell resolution, as well as the level of heterogeneity in the proteomic profiles of individual bacteria.

## RESULTS

### Sample Preparation Workflow Optimization

In this study we investigate how to push the sensitivity of SCP to a level where single bacteria can yield meaningful results. We thereby combined and adopted our robust and sensitive One-Pot^18^ SCP sample preparation workflow with the capability to isolate bacterial cells, making use of the microLIFE mode on the cellenONE, and the enhanced sensitivity of our LC-MS platform^4^ by employing columns with integrated emitter tip and zero dead volume, short 12.5 min active gradients, and the Orbitrap Astral MS. Our optimization process included screening gram-negative (*E. coli)* and gram-positive (*B. subitlis*) bacteria as well as staining and cell lysis conditions, including cell wall-deficient bacteria.

We started to tune sample preparation parameters for bacSCP using *E. coli* cells. We first aimed to directly apply our One-Pot workflow^18^, where cell lysis is enabled by the MS compatible detergent N-Dodecyl-β-D-maltoside (DDM) and incubation at 50 °C only, to bacterial cells. To do so, we removed the peptidoglycan cell wall prior to sorting to facilitate bacterial lysis by DDM. Bacteria were treated with Lysozyme to generate protoplasts or spheroplasts surrounded by a cell membrane only. The resulting cell-wall-deficient bacteria are spherical shaped and of < 2 μm diameter as estimated on the cellenONE. While the success of cell wall digestion for selected cells can be ensured by their altered shape on the cellenONE robot, their accurate isolation proved challenging as cells were often detected outside the PDC capillary by the image detection software of the cellenONE. This was likely caused by minimal movements of the PDC and dust or dirt particles by accident were detected as a cell. Activation of the reduce-shaking option on the cellenONE helped but still did not yield satisfactory results. Manual validation of detected particles inside the capillary proved difficult since no clear cell shape was visible by eye on the recorded pictures. As a result, alternative cell isolation and lysis procedures were explored.

We subsequently investigated whether repeated freeze-thaw cycles followed by heating of intact bacteria lead to successful lysis as previously demonstrated by Végvári et al^10^. Furthermore, we tested the effect of inhibition of cell division using cephalexin prior to cell wall removal, resulting in enlarged spheroplast cells^19^ and improved detectability on the cellenONE (estimated diameter by cellenONE 6.7 +/-1.6 μm). Of note, we expect reduced physiological relevance for cephalexin enlarged cells, but decided to include this condition to explore size limits needed for accurate detection using our setup on the cellenONE robot. Both strategies enabled us to successfully sort cells into individual wells of a 384 well pate. Introduction of a staining step (HOECHST 33342) ahead of sorting with the resulting fluorescence signal used as additional selection criteria on the cellenONE allowed us to improve the rate of confident successful isolations close to 100%. An overview of those tested workflow variations is depicted in Figure 1.

**Figure 1:**
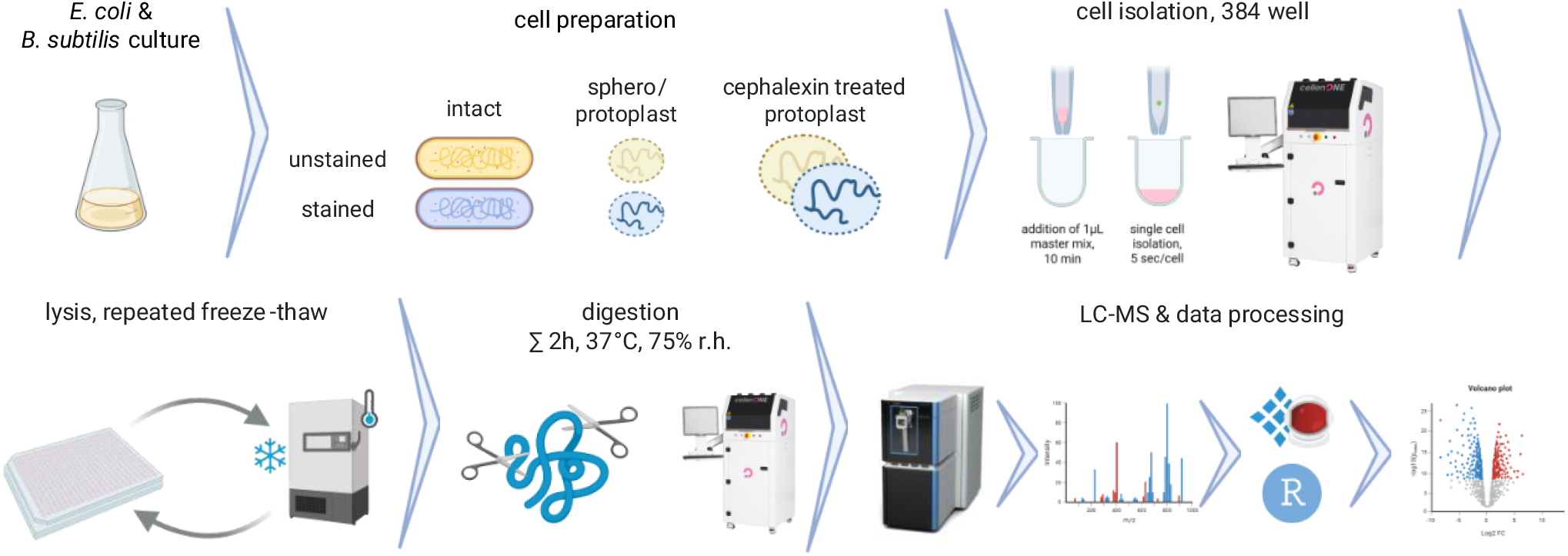
Workflow overview: To compare starting conditions, wild-type (intact) bacteria (stained or unstained) or cephalexin-enlarged protoplasts or spheroplasts were harvested at an OD600 of 0.7-0.9 followed by isolation of individual cells using the cellenONE robot. Cells were isolated into 1 μL of a pre-dispensed lysis and digestion mix within a 384 well plate. For intact cells, lysis was achieved by repeated freeze-thaw cycles. After stepwise digestion and a total of 2 h incubation, digestion was stopped by acidification using TFA and cells were directly subjected to LC-MS analysis in the very same 384 well plate using an 8 cm IonOpticks Gen4 column at 80 samples per day (SPD) throughput and an Orbitrap Astral MS. Figure created with biorender.com.

We first compared the above-mentioned cell-isolation and lysis conditions using *E. coli* cells and found 34 to 67 *E. coli* proteins on average per cell from intact unstained or stained cells respectively. The additional enlargement of spheroplasts using cephalexin yielded 92 *E. coli* proteins per cell on average, given that these spheroplasts originate from a fusion of several division-inhibited *E. coli* cells. On a limited number of replicates, we further analyzed 10 cell mini-bulks yielding 236 bacterial proteins on average but surprisingly no detectable effect on the number of quantified proteins was originating from cephalexin treatment (Figure 2A). We included the CRAPome^20^ database to account for contaminants, mainly originating from added trypsin as well as human skin keratins within our samples. Excluding those contaminants, we plotted the quantitative distribution of all proteins seen in 200 pg diluted *E. coli* bulk digests and matched the proteins found in single intact stained *E. coli* to the quantities from the bulk analysis. In line with our expectation, we were able to predominantly quantify higher abundant proteins from single cells within this distribution (Figure 2B). Of note, our 200 pg reference bulk already correlates to a low-input proteomic sample with input amounts comparable to single mammalian cells (i.e. HeLa cells).

**Figure 2:**
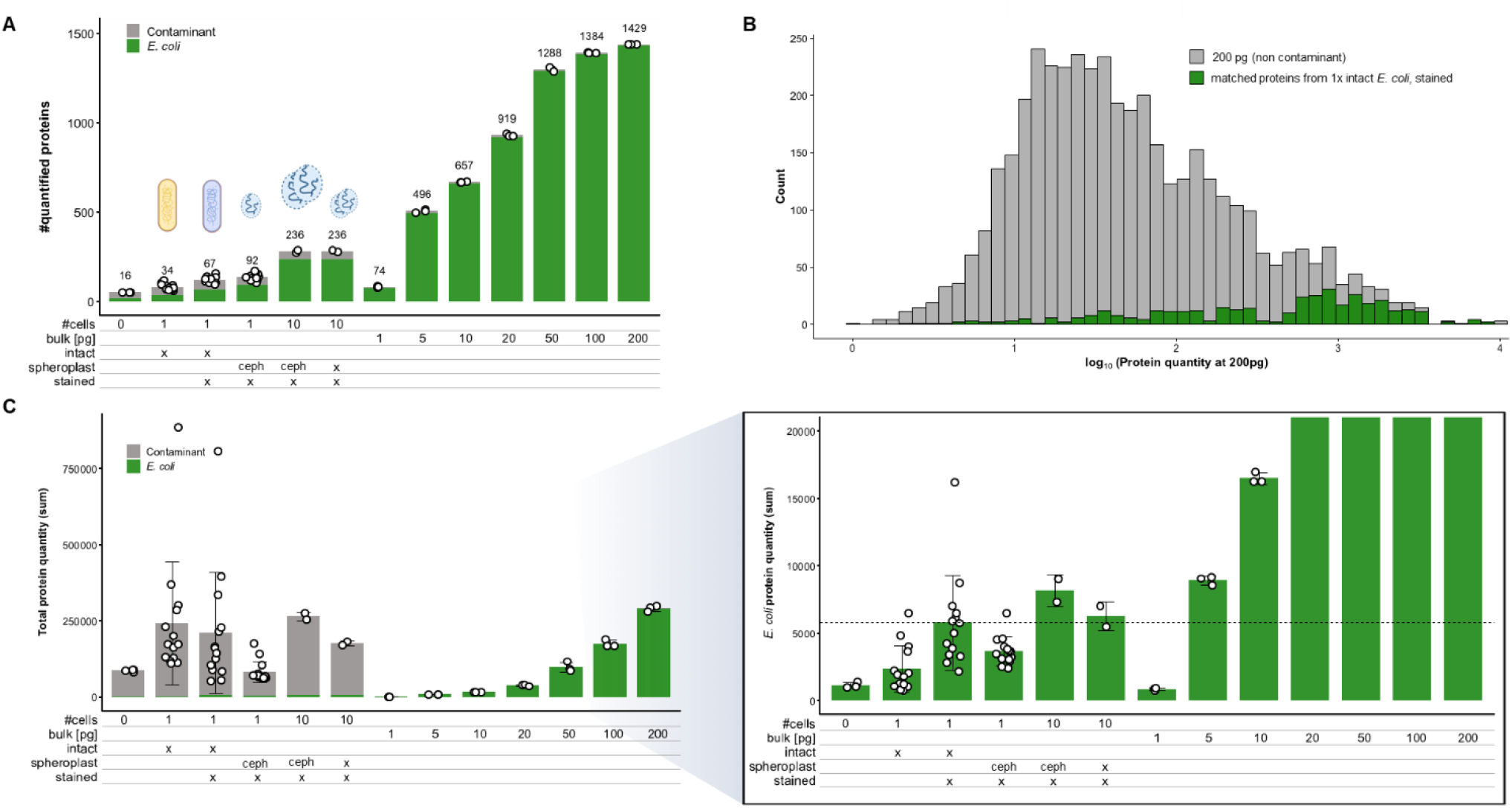
Establishing BacSCP using E. coli: **A**: Bars indicate the average number of quantified protein groups per single cell, spheroplast, 10 cells or from diluted bulks as indicated. White circles indicate the number of quantified proteins from individual replicates. Numbers above bars indicate the number of quantified non-contaminant proteins. **B.** Histogram showing the distribution of protein quantities found from 200 pg diluted bulk and their matched quantities from all n=14 measured single cell replicates of stained intact E. coli. **C**: Bars show the average summed protein quantity per cell, 10 cells or diluted bulks as indicated (left), as well as a zoom in with quantities from contaminant proteins excluded (right). Error bars show their standard deviation.

To estimate the relative total abundance of all quantified proteins in single cells or from 10 cells we compared the summed protein quantity to what was found from a titration of diluted bulk digests ranging from 1 pg to 1000 pg (Figure 1C). In line with previous reports^10^ and in line with our expectations (as a considerable excess of Trypsin protease was added to each sample) up to 98% of the total quantity of all proteins seen from single cells originated from contaminations, which is expected given the anticipated protein content from a single bacteria i.e. compared to the amount of protease added. Excluding those contaminants allows for improved comparison to the mini-bulks and an estimated 1 – 5 pg protein content extracted from single bacteria, or ∼5 pg from 10x bacteria respectively. Assuming that the 10× cell mini-bulk sample might deliver more accurate estimates, we assume having <1 pg protein/*E. coli*. This is in the same range as estimated by Lenski et al, who reported 865 fg per *E. coli*^9^ when in late exponential phase, as is the case for our cells.

To analyze the distribution of molecular processes associated with our single cell protein hits compared to bulk data, we mapped gene ontology (GO) slim terms to the respective protein hits (Supplementary Figure 1A). In bulk data, proteins involved in primary metabolic processes constitute the major fraction of annotated proteins, while in single cell protein hits this is shifted in favor of proteins belonging to the translation group reflecting the high abundance of ribosomes in the cell^21^ and the fact that we detect predominantly highly abundant proteins from single bacteria (Figure 2B). For stained single cells (intact or cephalexin-enlarged spheroplasts) we detect a higher amount of GO biological process groups, likely due to a higher number of absolute protein hits (Figure 2A). Mapping the GO cellular compartments (Supplementary Figure 1B), we observe a slight overrepresentation in plasma membrane associated proteins in the cephalexin-treated sample, potentially reflecting a higher membrane fraction or dysregulation of membrane components due to antibiotic treatment.

Based on the success of isolation confirmed by fluorescence and the wish to stay as close as possible to native conditions prior to lysis, we decided for intact stained cells as preferred condition. We therefore investigated the performance of our bacSCP workflow using gram-positive *B. subtilis* bacteria and the most promising conditions based on the previously recorded *E. coli* data. We thereby focused on intact cells and protoplasts to see potential effects on lysis success. We quantified ∼36 bacterial proteins on average/cell and ∼95 proteins/10 cells independent of the sample preparation strategy (Figure 3A) and a comparable ratio of bacterial to contaminant originating total protein quantity as seen for *E. coli* (Figure 3B).

**Figure 3:**
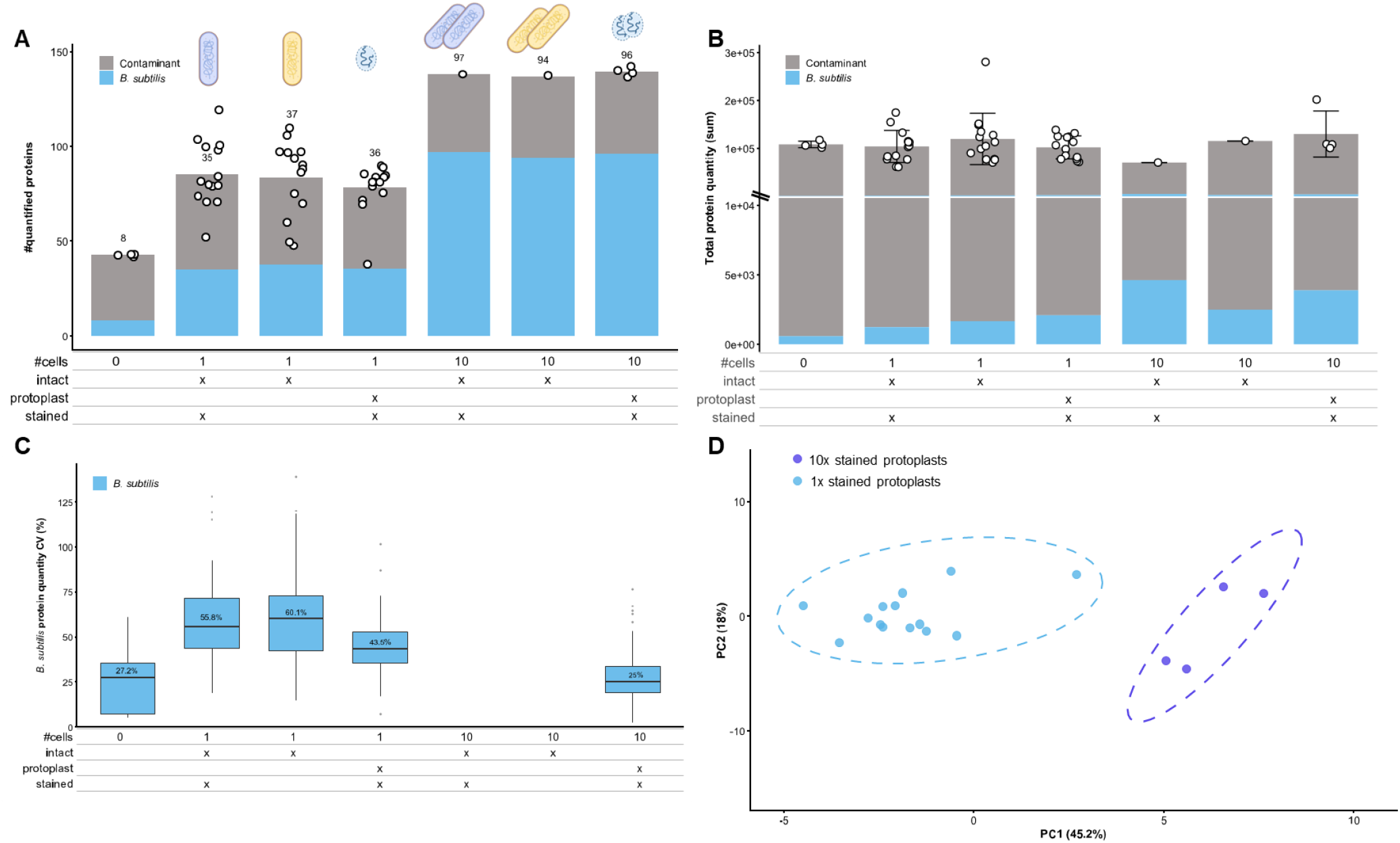
Setup of BacSCP using B. subtilis: Bars indicate the average number of quantified protein groups (**A)** or the average summed protein quantity (**B**) per single cell, protoplast or 10 cells as indicated. White circles indicate results from individual replicates. Numbers above bars indicate the number of quantified non-contaminant proteins. Error bars show standard deviations. **C**: The Boxplot shows the coefficient of variation of protein group quantities for all non-contaminant protein quantified in at least n=3 replicates, intact 10x mini-bulks were measured only once which is why no CV can be calculated there. **D**: Principal Component Analysis from all n=14 measured stained single protoplasts and n=4 mini-bulks from stained 10x protoplasts.

Coefficients of variation (CV) from bacterial protein quantities rise to ∼60 % on single cell level and 25% for 10 cells respectively (Figure 3C). While this seems high at first sight, it fits expectations as single-cell measurements using fluorescent protein reporters reported substantial cell-to-cell variability in protein expression, with a CV of ∼0.4.^22^ While technical variability for 250 pg inputs was shown to be below 10% using our setup^4^, we assume that for substantially lowered inputs as given on single bacterial level, technical variability can easily reach levels of 20% and more as seen on 10x cell level in Figure 3C, summing up to a total CV of ∼60% from biological and technical noise. A higher technical noise level for the majority of quantified precursors and proteins is to be expected based on a median of only 2 datapoints sampled by the MS per elution peak per precursor and only 2 precursors on median used for quantification per protein (Supplemental Figure 2). Of note, some precursors yielded much higher numbers of up to 20 datapoints per elution peak, likely yielding an improved quality of quantification. We were further wondering why the CV for the zero-cell negative control seems lower compared to single cells, although no bacterial input is present. We hypothesize that mainly noise rather than signals from peptides are recorded, and noise levels being on a more constant level (no biological variation). When including contaminants into the CV calculation, which introduces real peptide signals to that sample condition, the median CV drops a bit (from 27 to 25 %, Supplemental Figure 2C). Filtering for quantified proteins based on at least 3 precursors removes all hits of the 0 cell negative control, confirming no confidently quantified bacterial proteome hit hidden in the negative control (Supplemental Figure 2D). This very same filter helped to reduce the median CV of stained intact single cell samples from ∼56% down to 32%, presumably improving quantification accuracy.

Principial Component Analysis (PCA) of single cells versus 10 cells shows a clear separation of sample groups (Figure 3D), proving that quantitative differentiation of single to multiple cells works highly sufficient using a label free workflow. Together with the differences in summed quantity shown in Figure 3B, this confirms earlier results by Végvári et al.^10^ who was using an isobaric multiplexed approach and a carrier proteome to boost sensitivity on an Orbitrap Fusion Eclipse MS in 2023.

### Estimation of needed sensitivity level for finding biologically relevant protein regulation

Applying the now established workflow for bacSCP we next aimed for validation if a biologically relevant regulation of proteins could be observed at single bacteria level. To this end, we used a heat-stressed sample to investigate how much protein input could be reduced while still detecting the upregulation of the critical heat stress chaperones GroEL-GroES and ClpC. In *B. subtilis*, the ClpCP complex degrades proteins marked by the unusual post-translational modification phospho-arginine (pArg)^23^ and pArg-dependent degradation is critical for survival after prolonged heat shock^23^. Notably, phosphorylation of protein arginine residues is performed by the protein arginine kinase McsB and counteracted by the pArg phosphatase YwlE^24–26^. Here, we selected a *B. subtilis ΔywlE* strain, which is deficient in the pArg phosphatase YwlE and an established strain for detecting pArg-dependent effects^17,26^.

To analyze if the sensitivity given by our existing LC-MS setting is sufficient to accurately quantify relevant heat shock proteins we first acquired data from diluted bulk samples ranging from 1 – 1000 pg total input. We find that the regulation of groEL, groES and clpC upon heat stress is visible and keeps being significant for inputs down to 50 pg (Figure 4A-C). For very low inputs of 10 pg and less, data quality starts to drop (Figure 4D-F) and becomes sparser. At 1 pg injection amount those proteins were below their limit of quantification at 37 °C, while upon heat stress groES and groEL are quantified as evident in the ranked proteins (Figure 4G,H).

**Figure 4:**
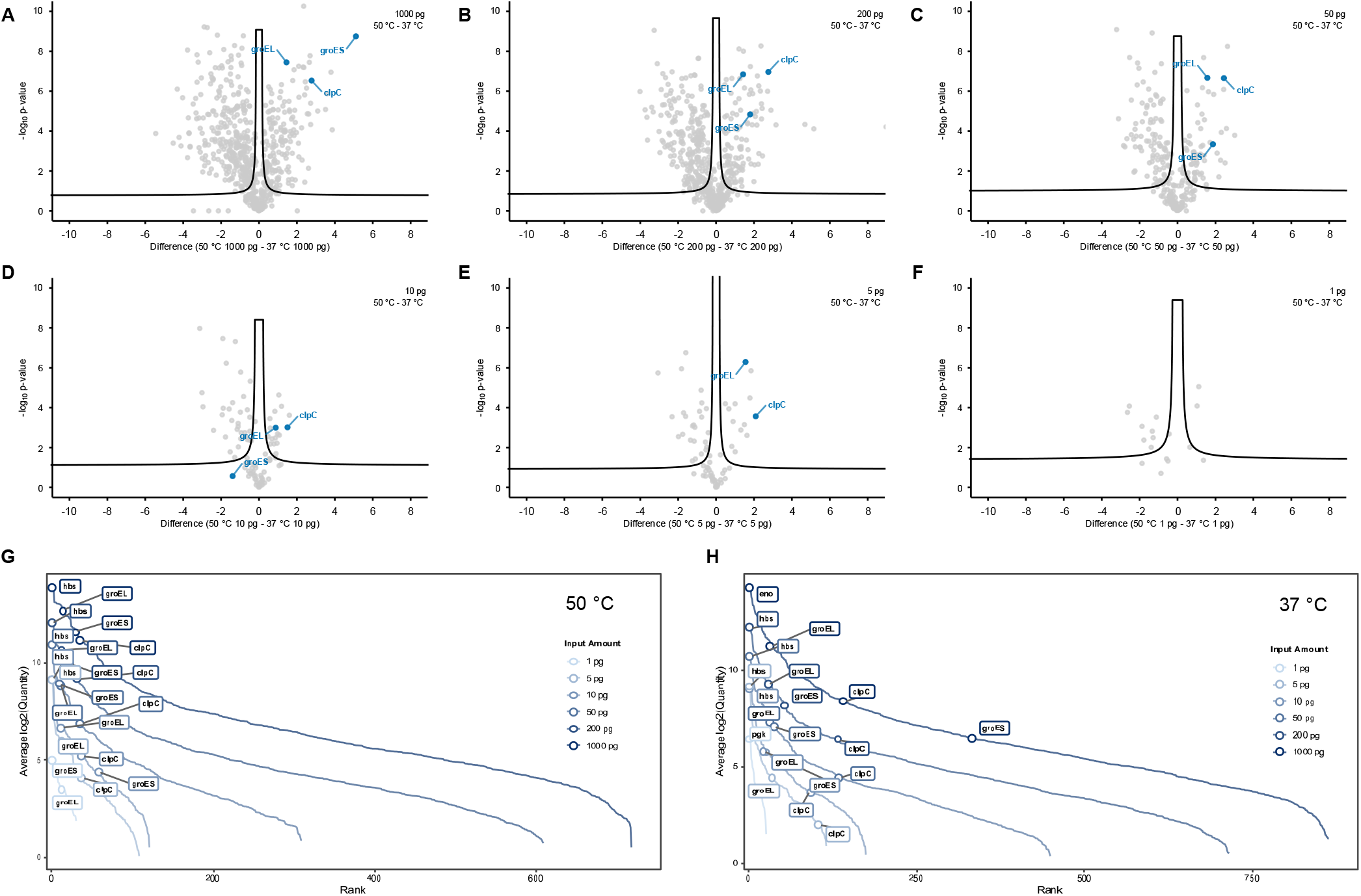
Sensitivity titration using B. subtilis ΔywlE bulk samples: **A-F**: Volcano plots from 1-1000 pg diluted bulk samples from heat shocked (50 °C) vs non heat shocked (37 °C) B. subtilis. GroEL, GroES and ClpC are highlighted. n=3 technical replicates, no imputation applied **G**: Ranked protein groups by their average quantity from those n=3 replicates for heat shocked (left) or non-heat shocked (right) samples, with the most abundant protein and proteins as in A-F highlighted.

### Heat shock response on *B. subtilis* wild type and mutants

Affirmed by these results, suggesting a potentially sufficient sensitivity level for quantifying chaperones in heat shock response, we continued benchmarking this regulation in different *B. subtilis* mutants, aiming to select the best cell line for a later single cell level experiment. We tested *B. subitilis ΔywlE* and *ΔmcsB* and the wild-type strain (Figure 5). Both *ΔywlE* and *ΔmcsB* are disrupted in pArg-dependent regulation, as either the pArg phosphatase YwlE or the pArg kinase McsB are missing, resulting in substantially increased pArg levels or complete lack thereof. ClpC as well as GroEL were shown previously to be modified by pArg and upregulated upon YwlE knockout^17,26^. Here, ClpC, GroEL and GroES are significantly upregulated during heat shock in all conditions tested (Figure 5). While in *B. subitilis ΔmcsB* the fold change of upregulation is lower than for wild-type or *ΔywlE* (Figure 5D), ClpC ranks overall among the top 10 most abundant proteins.

**Figure 5:**
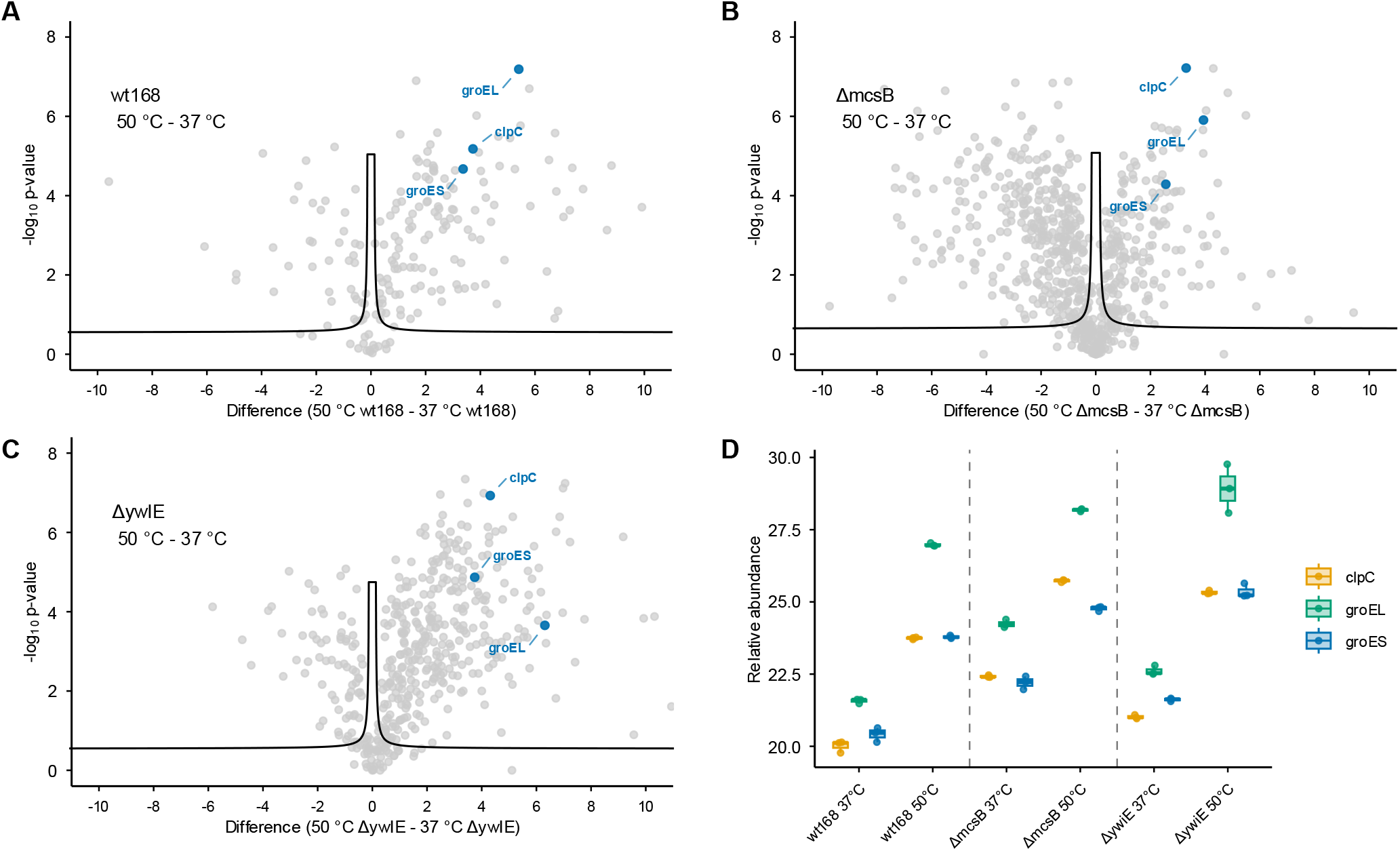
Comparison of B. subtilis variants. **A-C**: Volcano plots from 200 ng bulk samples from heat shocked (50 °C) vs non heat shocked (37 °C) wt168 (**A**), ΔmcsB (**B**) or ΔywIE (**C**) B. subtilis with relevant chaperons expected to be upregulated upon heat-shock highlighted. n=3 technical replicates **D**: The boxplot indicates the distribution of relative protein abundance of those highlighted proteins after label free quantification.

### BacSCP is sufficiently sensitive and robust to detect heat shock response on single bacteria level

Finally, we applied our optimized bacSCP conditions to investigate if heat shock response can be confidently quantified on single bacteria level and if cellular heterogeneity would become detectable. For this proof of principle study, we selected *B. subtilis ΔmcsB* as this cell line showed the highest relative abundance of ClpC in our pilot experiment (Figure 5). We analyzed 20 bacteria for each condition, harvested at 37 °C or after a 50 °C heat shock treatment. Cells were sorted intact and stained, yielding 42 and 55 bacterial proteins quantified/cell on average for non-heat-shocked and heat-shocked cells respectively (Figure 6A). In line with our observations in *E. coli*, those proteins predominantly represent abundant proteins found from reference bulk proteomic runs (Supplemental Figure 3). All three investigated chaperones showed a significant upregulation in a volcano plot when comparing average quantities from all n=20 single cells measured per condition (Figure 6B) and their regulation also becomes obvious by their shift in a ranking of all quantified proteins (Figure 6E). A slight effect was further observed for pgk and ctc, with ctc only quantified after heat-shock. The ribosomal protein Ctc is known as general stress response protein that associates to ribosomes under stress^27^, while Pgk is an essential glycolytic phosphoglycerate kinase and both proteins were found to be upregulated during heat stress in wild type *B. subtilis*^28^.

**Figure 6:**
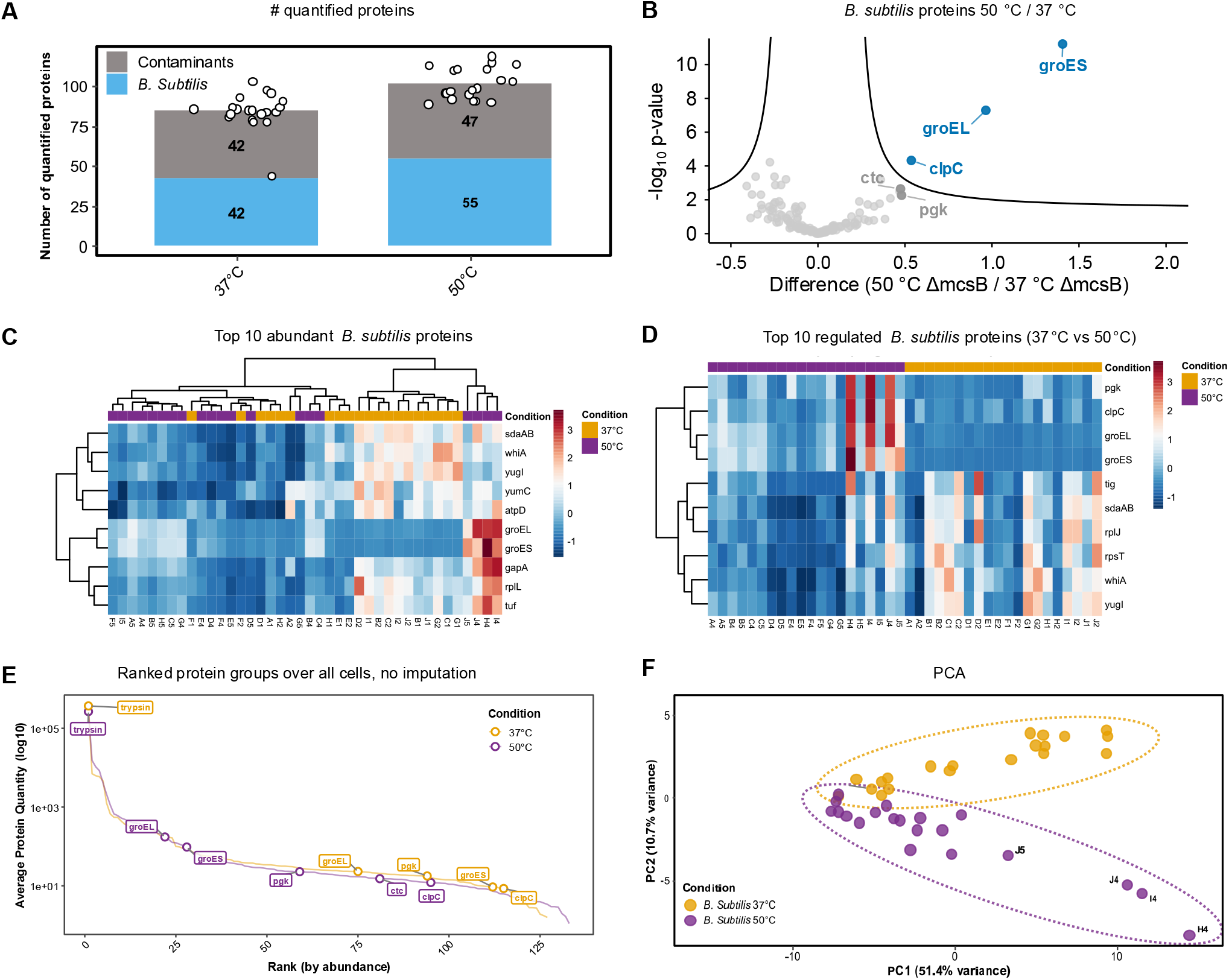
Investigation of heat shock response on single B. subtilis ΔmcsB: **A**: Bars indicate the average number of quantified protein groups from single B. subtilis with (50 °C) or without (37 °C) heat-shock, n=20 individual cells per condition. **B**: Volcano plot showing regulated non-contaminant proteins upon heat-shock. **C-D** Heatmaps showing relative quantities of the top 10 abundant or top 10 upon heat-shock regulated proteins. **E**: By quantity ranked protein groups found in all n=20 single cells measured/condition with most regulated proteins as seen in B highlighted. **F**: Principal component analysis from heat-shocked (50 °C) and non-heat-shocked (37 °C) single B. subtilis measured.

To investigate potential heterogeneity within the analyzed cell population, we further plotted a heat map of the top abundant as well as top regulated proteins (Figure 6C,D). We found a well-defined subset of proteins mainly upregulated upon heat-shock, with groEL and groES among the strongest upregulated candidates. A list of those regulated proteins including their biological function and reported pArg modification sites is shown in Figure 6. As observed for *E. coli*, the major biological processes involved are translation and metabolic enzymes, now joined by protein quality control factors induced by heat shock. Furthermore, a subset of 4 cells seems to show a much stronger heat-shock response and by that defines a separate cluster (Figure 3C). Considering high technical run-to-run variability expected for such ultra-low input proteomic samples (see Figure 3C), we assume that the visible heterogeneity might at least partly result from technical variance and partly from biological heterogeneity. A significantly enlarged sample size would be needed in future studies to improve statistical power and confirm our hypothesis. Given that this is the first study ever showing biological regulation on single bacteria level, we are convinced that our data still gives valuable first hints towards heterogeneity present also in cultured bacterial cells. This picture is further mirrored when doing a PCA, that shows a well-defined separation of non-heat-shocked vs heat-shocked cells and those cells with strongest response in groEL, groES and other proteins as shown in the heatmap forming a distinct population also in the PCA (**Error! Reference source not found**.F).

**Table 1:**
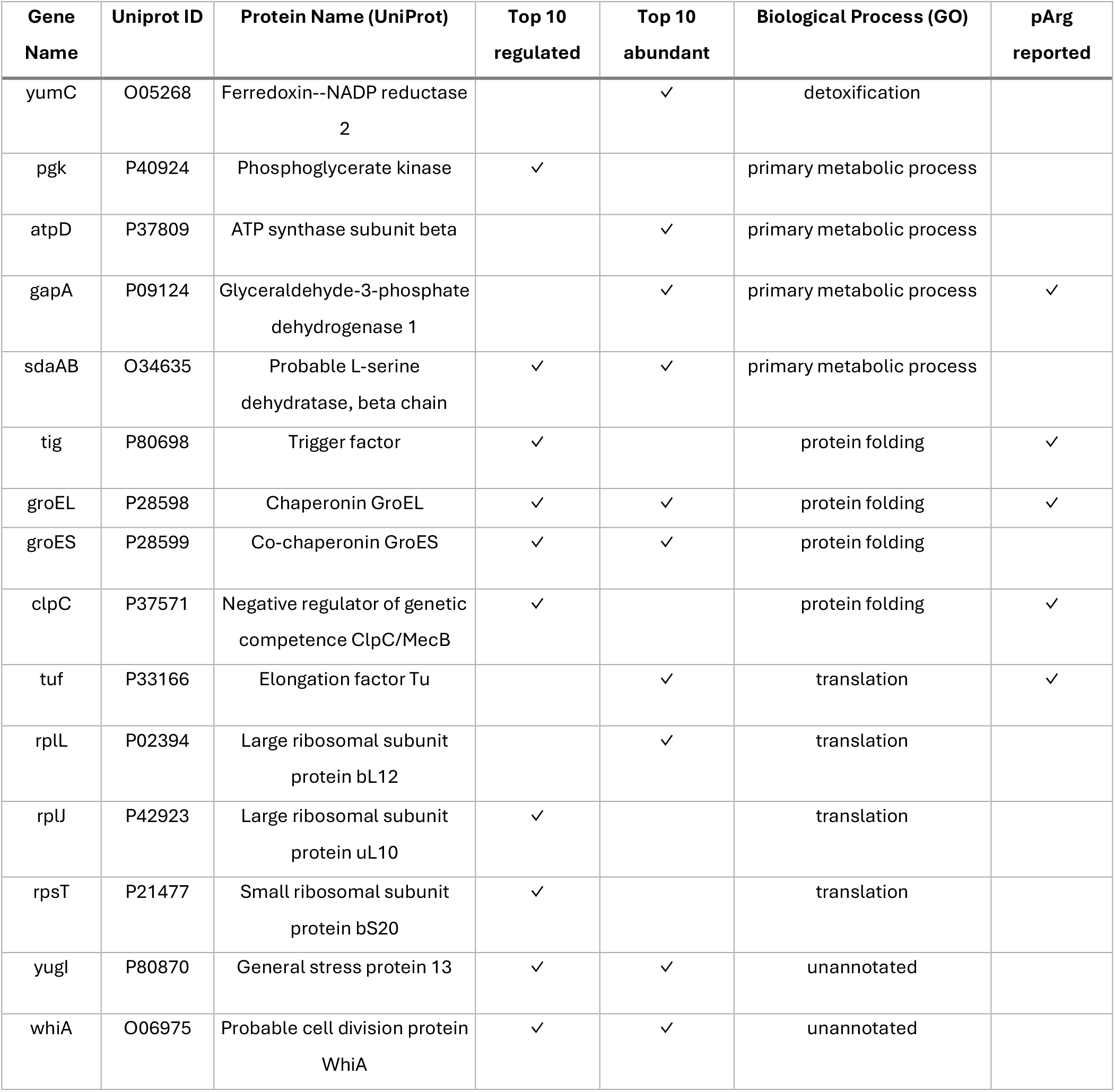
Top10 regulated and Top10 abundant proteins from Figure 6C and D. Biological Process GO slim data was retrieved from the QuickGo^29^ database, modification by pArg from Schmid et al.^30^ and Elsholz et al.^25^.

## DISCUSSION

We demonstrate a bacterial single cell proteomics method effective for both the gram-positive (*B. subtilis)* and gram-negative model organisms (*E. coli)*, identifying a comparable number of non-contaminant proteins (∼ 55 vs 65 respectively). In accordance with cellular abundance, we predominantly detect metabolic enzymes and ribosomes, and in the case of heat-stressed *B. subtilis*, stress response chaperones. However, our technique also detects proteins in the medium abundance range (Figure 2B, Supplementary Figure 3). While accurate quantification in this range is currently unfeasible, it nevertheless highlights the sensitivity of our method. Of note, the only previous study published back in 2023^10^ used a multiplexed approach including a 250 cell carrier channel in order to boost sensitivity to a level where quantitative differentiation of single to multiple cells was possible. With current state of the art loss-less workflows and sensitive MS instrumentation, such a differentiation is possible by means of label free proteomics without the use of any library or carrier (Figure 3D).

Strikingly, we could furthermore detect the upregulation of GroEL, GroES and ClpC in heat-stressed *B. subtilis* samples at single cell level. This effect was most pronounced in four individual cells (Figure 6D). Protein levels of GroEL, GroES and ClpC are relatively uniform at 37 °C (more so for GroEL/ES than for ClpC) while protein levels after heat shock differ markedly. We observe three groups of single cells where the targeted proteins of interest are either barely, intermediately or highly enriched, indicating strong heterogeneity in response of individual cells to heat stress. Interestingly, GroEL/ES and ClpC are under the transcriptional control of different repressors, HrcA and CtsR respectively^31,32^, but in cells we observe concurrent increased protein abundance of ClpC, GroEL/ES as well as the not stress related protein Pgk, indicating an overall heat stress effect rather than effects specific to certain proteins or regulons. However, caution is required when interpreting these results, as due to the very low input sample and the high technical variability, some observed effects could be technical rather than biological in nature. Future experiments that extend beyond a proof-of-concept study will require a larger sample size to validate our results.

Expanding this approach to include additional bacterial species in future experiments would provide further insights into the generalizability of our results. In particular, the application of this method to pathogenic bacteria is of interest, if technically possible. In line with our data on ClpC, bacSCP analysis of *Staphylococcus aureus* where the ClpC operon via McsB is required for virulence^33^, or Mycobacteria (e.g. *Mycobacterium tuberculosis, Mycobacterium smegmatis*), where ClpC1 is a target of antimicrobial compounds like Cyclomarin or Ecumicin, might provide valuable insights into differential regulation of this important quality control factor^34,35^.

To further delve into pathogenic bacteria heterogeneity and proteins involved in antibiotic tolerance and resistance, bacSCP will still need to improve substantially in terms of sensitivity, as often pathologically relevant proteins are not among the most abundant proteins in the cell^36^. However, we anticipate that future technical and methodological improvements of bacSCP will further push the limit of detection. While analysis of single bacterial cells has been established for other techniques, such as phenotypic descriptions of cell processes via fluorescent reporters, transcriptome sequencing, metabolome analysis by MS^12,14^, this is to our knowledge the first attempt at bacterial single cell proteomics where changes to single cells upon stress were observed.

## METHODS

### Culturing bacterial cells

*B. subtilis ΔywlE* was generated as described in^30^ and *B. subtilis ΔmcsB* was generated as described in^37^. *Escherichia coli* strain BL21(DE3), *B. subtilis* strain 168, wild-type and *ΔywlE* and *ΔmcsB* knockout strains were inoculated from a glycerol stock into LB medium and grown overnight at 37 °C with constant agitation of 140-160 rpm. The main culture was inoculated with 1/100 of the overnight culture. At an OD_600_ of 0.7-0.9 samples were either harvested by centrifugation at 4 °C, or transferred to a preheated shaker and subjected to heat shock for 20 minutes at 50 °C and subsequently cooled down on ice and harvested. Cell pellets were washed 3x times with PBS, resuspended in Buffer 50 mM Hepes pH 7.3, 100 mM NaCl + 25% glycerol, flash frozen in liquid N_2_ and stored at -80 °C until analysis. Stained cells were incubated before analysis in a 1:1000 Hoechst 33342 (Thermo Scientific Pierce) dilution in PBS for 30 minutes at room temperature in the dark and washed 3x in PBS and diluted to ∼200 cells/μL prior to sample analysis.

### Generation of protoplast/spheroblast cells

Wild-type and cephalexin-enlarged *E. coli* spheroplasts were prepared as described in Figueroa et al^38^. In short, *E. coli* were grown as described in 25 mL LB to an OD_600_ of 0.7-0.9 and intact *E. coli* samples were harvested. To produce cephalexin-treated *E. coli* cells were diluted 1:10 in 30 mL LB containing 60 μg/mL cephalexin (Sigma Aldrich) and incubated at 37 °C for 2.5 hours. Filaments were harvested at 4 °C. To produce spheroplasts, cell pellets were washed with 0.8 M sucrose and incubated in 350 mM Tris-Cl pH 7.8, 1.5 mg/mL Lysozyme, 0.35 mg/mL DNAse and 35 mM EDTA pH 8 for 10 mins at room temperature. 1 mL solution A (20 mM MgCl2, 0.7 M sucrose, 10 mM Tris-Cl, pH 7.8) was added and cells incubated for 4 minutes. Subsequently, 7 mL of solution B (10 mM MgCl2, 0.8 M sucrose, 10 mM Tris-Cl, pH 7.8) were added at 4 °C and pellets gently harvested and resuspended in 700 μL Solution B.

*B. subtilis* protoplasts were prepared similar to Chang & Cohen^39^. *B. subtilis* were grown as described in 25 mL LB to and OD_600_ of 0.7-0.9 and harvested. Pellets were resuspended in 4 mL of SMM (0.5 M sucrose, 0.02 M maleic acid pH 6.5, 0.02 M MgCl2) supplemented with 2.5 mg/mL Lysozyme and incubated for 35 minutes at 37 °C under gentle agitation. Pellets were harvested and resuspended in 500 μL SMM. Success of cephalexin treatment and spheroplast/protoplast formation was confirmed by bright field microscopy. Aliquots of the suspension were flash frozen in liquid N_2_ and stored at -80 °C until analysis.

### Sample preparation for *B. subtilis* bulk samples

For bulk analysis, pellets of the respective *B. subtilis* strains were thawed in the presence of Benzonase endonuclease and 5 mM MgCl2 and lysed by bead beating (Precellys 24, Bertin Technologies). 1.5 mL screw cap Eppendorf tubes were filled with ∼500 μL of acid-washed glass beads <106 μm (Sigma Aldrich) and 700 μL of cell suspension and run for 6800 rpm, 3x 20 seconds, 30 seconds pause at 4 °C. Lysates were cleared by centrifugation at 4 °C. Lysate concentration was measured by BCA Assay (Pierce). Lysates were diluted to 1 mg/mL and digested using trypsin (Trypsin Gold, V5280, Promega, 500 ng per sample) over night at 37 °C. Digestion was stopped by acidification to a final concentration of 1% (v/v) TFA and peptide concentration was estimated by means of UV-VIS chromatography at 214 nm using a monolithic (PepSwift™ Monolithic 50 mm x 0.2 mm) using the following gradient at a constant flow of 1.5 μL/min: 3 min 2% eluent B (80% ACN, 0.04% TFA), increasing to 95% over 20 min and kept at 95% for 5 min, decreasing to 2% in 1 min before re-equilibrating at 2% for 6 min. Eluent A was 0.05% TFA in water.

### Sample preparation for single cell proteomic samples

Isolation of single intact *E. coli* or *B. subtilis* as well as their protoplasts/spheroplasts, cephalexin treated or stained cells was performed within the cellenONE X1 Neo robot. Thereby cells were sorted into a 384 well plate (Eppendorf twin-tec PCR LoBind) followed by lysis and digestion based on our previously published One-Pot protocol (Matzinger et al, Anal Chem, 2023) with the following details: Cells were sorted into wells containing 1 μL of master mix (0.2% DDM (D4641-500MG, Sigma-Aldrich), 100 mM triethylammonium bicarbonate (TEAB; 17902-500ML, Fluka Analytical), 3 ng μL^–1^ trypsin (Trypsin Gold, V5280, Promega) and 1% DMSO). Humidity and temperature were controlled at 30% and 10 °C during cell sorting. Cells were isolated in microLIFE mode and stained cells were furthermore sorted in the basic cellenONE mode using the following parameters: Allowed size 4-10 μm (sizes starting at 1 μm for microLIFE mode) at a maximum elongation of 2, for stained cells with blue fluorescence channel active at 40% gain and 40 ms exposure time, selection with a minimum fluorescence intensity of 10. Protoplasts and spheroplasts were directly heated with lysis happening in parallel to digestion. For intact cells, lysis was facilitated by 5 freeze thaw cycles of at least 30 min freezing time at -70 °C and subsequent thawing at room temperature for 15 min each and protein digestion was performed at 50 °C and 80% relative humidity for 30 min before an additional 500 nL of 3 ng μL^–1^ trypsin was added followed by incubation for final 1.5 h at 37 °C and 70% relative humidity. Digestion was stopped by addition of 2 μL 0.5% (v/v) TFA and storage at −20 °C until MS analysis. For LC–MS/MS analysis, samples were directly injected from the 384-well plate.

### LC–MS analysis

Samples were analyzed using the Thermo Scientific Vanquish Neo UHPLC system. Peptides were separated on an Aurora Rapid 8x75 XT Gen4 nanoflow UHPLC column with an integrated emitter (IonOpticks) at 50 °C using a NanoShield C18 prototype trap column (IonOpticks).

Peptide separation was performed at 80 SPD with the following details: For the first 0.5 min a flow rate of 1 μL/min at 12% buffer B (80% ACN, 19.9% ddH2O, 0.1% TFA) was applied. Until minute 0.6 the flow rate was reduced to 250 nL/min at 13% buffer B followed by a stepwise gradient to 22% buffer B until minute 3, 37% buffer B until minute 10.5 and 56% buffer B until minute 12.5 respectively. After that the flow rate increased to 1 μL/min at 99% buffer B until minute 13 for column washing and was kept in this state for 1 min until minute 14. Fast sample loading was performed at a maximum pressure of 800 bar and at a maximum flow rate of 10 μL/min.

For MS measuring, the Orbitrap Astral MS (Thermo Scientific) equipped with a FAIMS Pro Duo interface (Thermo Scientific) and an EASY-Spray source was coupled to the LC. FAIMS was used in low resolution mode^40^ with an outer electrode temperature of 80 °C and an inner electrode temperature of 100 °C at a compensation voltage of – 48 V. MS1 spectra were recorded using the Orbitrap analyzer at a resolution of 240,000 from m/z 400 to 800 using an automated gate control (AGC) target of 500% and a maximum injection time of 100 ms. For MS2 in DIA mode using the Astral analyzer, non-overlapping isolation windows of m/z 20 and a scan range from an m/z of 400 to 800 was chosen. A precursor accumulation time of 60 ms was allowed for and the AGC target was set to 800%.

### Data analysis

All raw data were analyzed using Spectronaut (version 20.4.260109.92449, Biognosys^41^). Data was searched library free using DirectDIA+ with replicates defined as the same condition in the search settings. Quantification was performed on MS1 level. Carbamidomethylation of cysteines as a static modification was removed as no alkylation step was performed with factory settings used besides that. Searches were performed against the *E. coli* K12 proteome (uniport reference proteome Tax ID 83333 as of 2025-06-30, 4402 sequences) or *B. subtilis* strain 168 proteome (uniport reference proteome, Tax ID 224308, as of 2025-06-27, 4194 sequences) respectively with the CRAPome^20^ (118 protein sequence entries) added to all searches as contaminant database.

### Post-processing

Statistical analysis and visualization were done in Perseus (v2.0.11)^42^ or using custom scripts in R (v4.5.2) within R Studio (v2026.01.0.392) or Python. Troubleshooting and finetuning of custom scripts was supported by AI based LLM using Abacus (abacus.ai). Workflow charts were generated using biorender. GO slim annotations were procured via the QuickGO browser (https://www.ebi.ac.uk/QuickGO/slimming)^29^, using the goslim_prokaryote_ribbon preset. Only hits assigned by Uniprot were taken, and redundant annotations removed.

## ACKNOWLEDGMENTS

This work was supported by the infrastructure funding 4th call 2022/01 (AT-SCP) of the Austrian Research Promotion Agency (FFG) as well as by the FWF projects 10.55776/PAT4800425 and 10.55776/PAT2059025. J.L. is funded by ESP 467 (Grant-DOI 10.55776/ESP467) of the Austrian Science Fund (FWF). All LC-MS/MS analyses in Vienna were performed on the Vienna BioCenter Core Facilities instrument pool. We would like to thank Tim Clausen for constant scientific support and providing feedback to the manuscript, members of the Clausen lab and Rupert Mayer for proofreading the manuscript. For the purpose of open access, the authors have applied a CC BY public copyright license to any Author Accepted Manuscript version arising from this submission.

## AUTHOR CONTRIBUTIONS STATEMENT

JL and MM conceptualized the study, designed, and performed experiments, data analysis, and wrote the manuscript. TT performed experiments and helped with post-processing. All authors revised and agreed on the manuscript.

