## Supplemental Figures for "Pushing the limits of SCP: bacSCP, a proof-of-concept study to investigate heterogeneity of bacteria by single cell proteomics"

### Content

Supplemental Figure 1: **Biological functions and cellular compartments of quantified proteins from E. coli**

Supplemental Figure 2: **Influence of precursors per protein and datapoints across peak on quantification accuracy**

Supplemental Figure 3: **Protein quantity distribution from single B. subtilis matched to bulk level analysis**

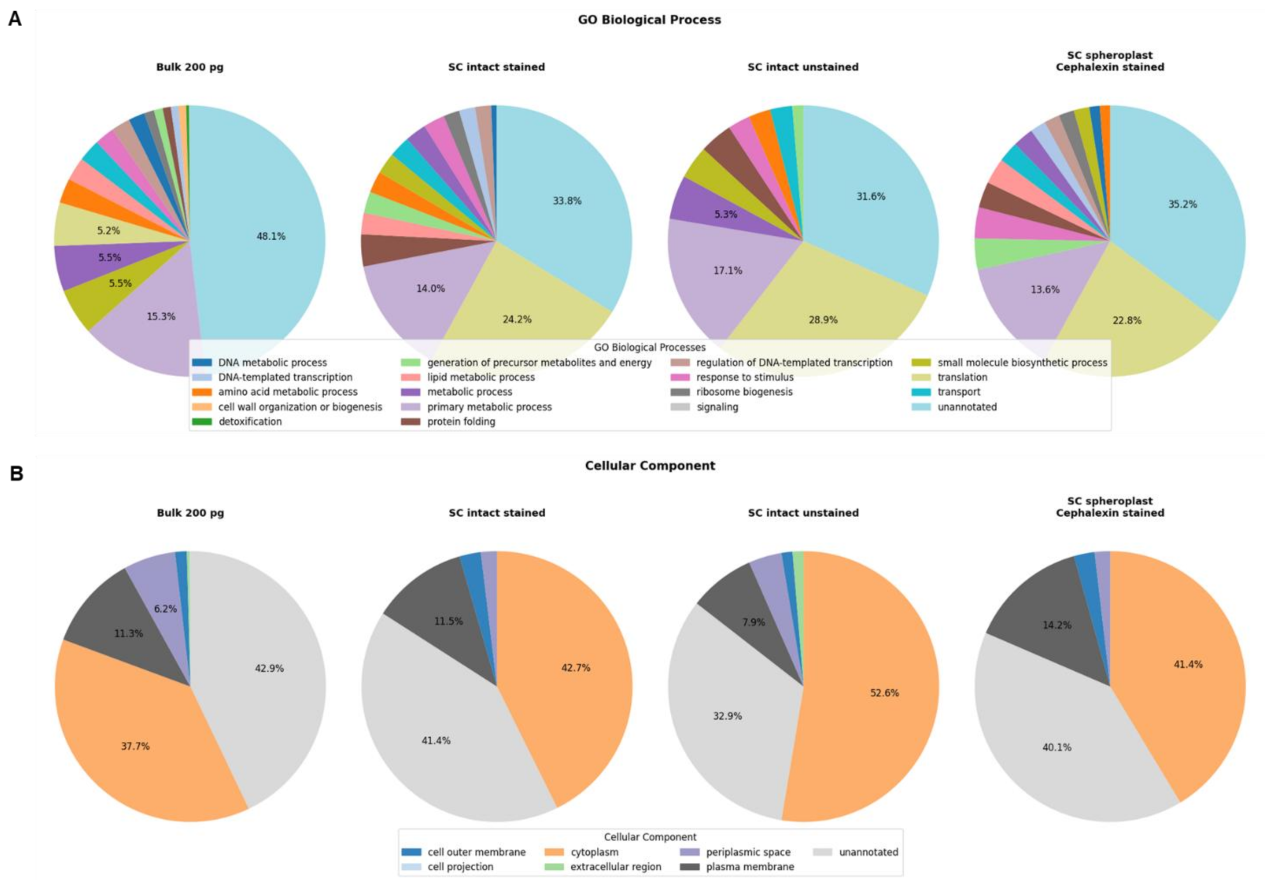


*Supplemental Figure 4:* ***Biological functions and cellular compartments of quantified proteins from E. coli.*** *GoSlim analysis of diluted bulk and single E. coli in dependence of sample preparation strategy as indicated. SC: single cell. Pie charts depict the biological processes (**A**) and cellular compartments (**B**) reported for those proteins.*

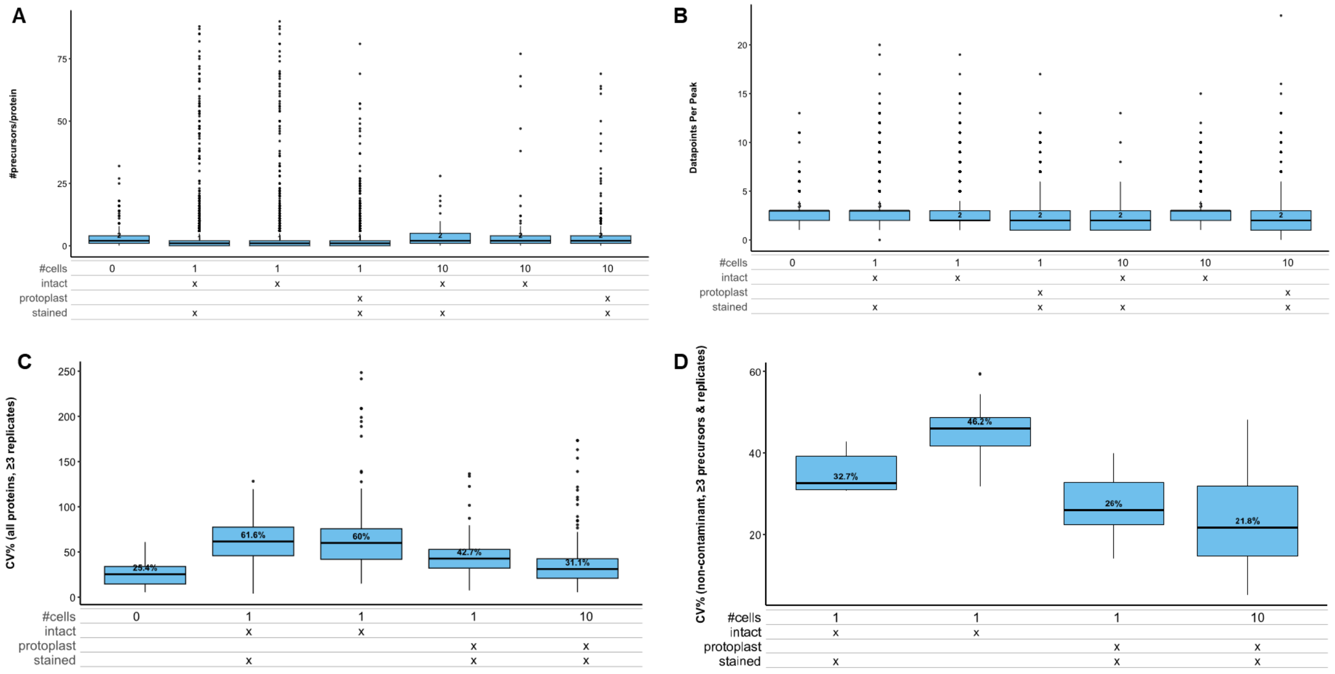


*Supplemental Figure 5:* ***Influence of precursors per protein and datapoints across peak on quantification accuracy*** *Data of Figure 3 is shown, the box-plots depict the distribution of number of precursors used for quantification per protein (**A**), the number of datapoints across the chromatographic elution peak used to quantify precursors on MS1 level (**B**), the CV on protein level including all proteins found in at least 3 replicates (**C**), or the CV of all bacterial proteins, excluding contaminants, found with at least 3 precursors and in 3 replicates (**D**).*

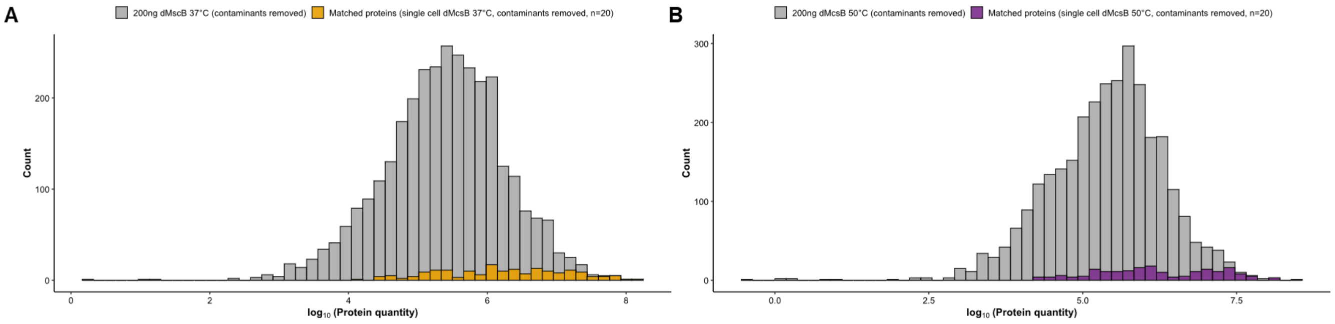


*Supplemental Figure 6:* ***Protein quantity distribution from single B. subtilis matched to bulk level analysis****. Histogram showing distribution of proteins found from 200 ng ΔmcsB B. subtilis and those from single cells matched to bulk quantities. Data shown for non-heat shocked, 37 °C (**A**) and heat shocked, 50 °C cells (**B**), respectively.*
